# Neural correlates of hand-object congruency effects during action planning

**DOI:** 10.1101/2020.10.16.342147

**Authors:** Zuo Zhang, Natalie Nelissen, Peter Zeidman, Nicola Filippini, Jörn Diedrichsen, Stefania Bracci, Karl Friston, Elisabeth Rounis

## Abstract

Selecting hand actions to manipulate an object is affected both by perceptual factors and by action goals. Affordances are associated with the automatic potentiation of motor representations to an object, independent of the goal of the actor. In previous studies, we have demonstrated an influence of the congruency between hand and object orientations on response times when reaching to turn an object, such as a cup. In this study, we investigated how the representation of hand postures triggered by planning to turn a cup were influenced by this congruency effect, in an fMRI scanning environment. Healthy participants were asked to reach and turn a real cup that was placed in front of them either in an upright orientation or upside down. They were instructed to use a hand orientation that was either congruent or incongruent with the cup orientation. As expected, the motor responses were faster when the hand and cup orientations were congruent. There was increased activity in a network of brain regions involving object-directed actions during action planning, which included bilateral primary and extrastriate visual, medial and superior temporal areas, as well as superior parietal, primary motor and premotor areas in the left hemisphere. Specific activation of the dorsal premotor cortex (PMd) was associated with hand-object orientation congruency during planning, and prior to any action taking place. Activity in that area and its connectivity with the lateral occipito-temporal cortex (LOTC) increased when planning incongruent actions. The increased activity in premotor areas in trials where the orientation of the hand was incongruent to that of the object suggests a role in eliciting competing representations specified by hand postures in LOTC.

## 1. Introduction

Gibson introduced the term ‘affordance’ (1979) to describe the context-specific influence of object properties on action goals. Affordances can elicit stimulus-response compatibility effects based on a correspondence between the graspable features of an object and an independent task-related action (Craighero et al. 1996, Castiello 1999, Creem and Proffitt 2001, Gentilucci 1998, 2002, Bub and Masson 2010). In a classical experiment, Tucker and Ellis (1998) demonstrated that the speed of finger press responses for object categorisation was influenced by the compatibility between the object orientation and hand response, even though participants were not required to make a judgement about the object orientation. Since then, a variety of perceptual tasks have shown that visual properties of objects can give rise to action representations (Chao and Martin 2000, Grezes and Decety 2002, Mahon et al. 2007). These effects are context specific and stronger with real objects (Gomez et al. 2017). In a study by Creem and Proffitt (2001), participants were observed to grasp objects by their functional side (eg. their handle, in the case of a saucepan), when performing a dual-visuospatial task, but not when performing a dual-semantic task, suggesting that affordances may elicit conceptual knowledge about objects rather than simple visuospatial mappings (Creem and Proffitt, 2001, Bub et al. 2018).

The elicitation of movement representations by objects is of fundamental importance in understanding higher order motor deficits in patients, described in the neuropsychology literature. Patients with a condition known as ‘alien limb syndrome’ (Riddoch et al. 1998; McBride et al. 2013), show movement-specific interference effects elicited by graspable features of objects (Riddoch et al. 1998). In another disorder, known as limb apraxia, there are deficits in exerting cognitive control over competing movement plans elicited by affordances (Rounis and Humphreys, 2015). These patients often demonstrate an over-reliance on familiar movements elicited by object affordances, at the expense of movements needed to complete the task goal (Lee et al. 2014, Watson and Buxbaum 2014, Pizzamiglio et al. 2020).

Another influence in movement selection that determines choice of trajectories when manipulating an object is the task goal (Bernstein 1967, Wolpert 1997, Harris and Wolpert 1998). One example, the ‘end state comfort’ effect, describes the preference for participants to start an action uncomfortably with a plan to use an intrinsically familiar trajectory to achieve an action goal, leading to a comfortable posture at the end (Rosenbaum et al. 1990, 1992). However, recent experiments have demonstrated situations where affordance effects trump the end state comfort effect. When participants are asked to turn a cup from its upright position, upside-down, they often favour a hand posture that is compatible with the object orientation and typically grasp it from its top (or open end), even though this would lead an uncomfortable posture in the end (Herbort and Butz 2011). This is corroborated by evidence that their reactions times are shorter in conditions where the hand and cup orientation are congruent, compared to when they are incongruent, when tested in a forced choice task (Rounis et al. 2017). In this situation, affordances, demonstrated by an ‘initial grasp preference’ in which there is congruency between the initial grasp posture and cup orientation, override the ‘end state comfort’ effect (Herbort and Butz 2011, Rounis et al. 2017).

The interplay between affordances and end-state comfort effects (ie. initial and end posture preferences) when moving an object (Drapati and Sirigu, 2006) is likely to be mediated by separable neural substrates (Owen 1997, Dickinson and Balleine, 1998, Packard and Knowlton 2002, Waszak et al. 2005, Herbort and Butz 2011, Rounis et al. 2017, Pizzamiglio et al. 2020), involving brain areas responsible for motor control, located in the dorsal stream (Rizzolatti and Mattelli 2003, Nachev et al. 2008, Wolpe et al. 2020) and action semantics, located in the ventral stream (Mahon et al. 2007, van Elk et al. 2014). Some functional imaging studies have reported the neural correlates of actions directed to real objects in the scanner. These have mostly contrasted between different actions (Valyear et al. 2007, Gallivan et al. 2011, 2013), or between different objects (Grol et al. 2007, Mahon et al. 2007, Sakreida et al., 2016, Fabbri et al. 2014 and 2016). Very few functional imaging studies have investigated the neural correlates of congruency effects elicited by affordance in healthy volunteers (Grezes et al. 2003, Kumar et al. 2012), which is at odds with the extensive body of behavioural literature of this effect, mentioned above. Nevertheless those that have investigated this effect, report a prominent role of dorsal premotor cortex in selecting alternative movement plans ( Grezes et al. 2003, Cisek and Kalaska 2005).

In this study, we explored the neural underpinnings of hand-object congruency effects, when planning to move a cup within an fMRI environment. We converted a task, in which participants had to turn a cup from one orientation to another, from a previous behavioural study (Rounis et al., 2017), into an ‘object in the scanner’ fMRI task. A handle-less cup was placed either upright or up-side-down, for participants to turn either using a supinated (‘straight’) or a pronated (‘invert[ed]’) hand grasp. Based on our previous results, we expected to find that grasps in which the cup and hand orientation were congruent (ie. ‘afforded’) would be faster. At the neural level, we investigated regional brain activations during motor planning, to reveal how congruency between the hand and the cup influenced areas involved in object manipulation (Drapati and Sirigu 2006, Mahon et al. 2007, Grol et al. 2007, Verhagen et al 2008, Gallivan et al. 2011, 2013), prior to any movement taking place.

## 2. Materials and Methods

### 2.1 Participants

Twenty-seven healthy righted-handed volunteers were recruited to participate in this study (14 females, 13 males; mean age = 27.95 years; age range = 20-38 years). All participants had normal or corrected-to normal vision. Full written consent according to the declaration of Helsinki was obtained from all participants. The study was approved by Oxford University’s Central University Research Ethics Committee (MS-IDREC-C1-2015-097). Participants were compensated £10/hr or course credits for participating in the experiment. Data from two participants were discarded because technical issues caused the behavioural and timing data not to be recorded. The study procedures or analyses were not pre-registered prior to the research.

### 2.2 Experimental setup

Participants performed an instructed-delay cup-manipulation task while lying supine in the MRI scanner. In this task, the cup ‘manipulation’ involved the action of reaching to and turning the cup from an upright position upside-down, or vice versa. Previous studies implicate different brain regions associated with moving an object, as opposed to using it (Drapati and Sirigu 2006).

The standard mattress of the scanner bed was replaced by a thinner one, allowing participants to lie lower within the scanner bore so that they could comfortably bend their head to look at the object positioned on a custom-made Perspex platform in front of them. Their head was positioned inside a phased array receiver 12-channel MRI headcoil, which rested on a 15° wedge (Figure 1). Participants’ overall head tilt was 25° from supine, considering the width of the headcoil and padding provided, which lifted their head further inside it. This allowed for direct visualisation of the cup to be grasped and visual control of their hand movement.

**Figure 1:**
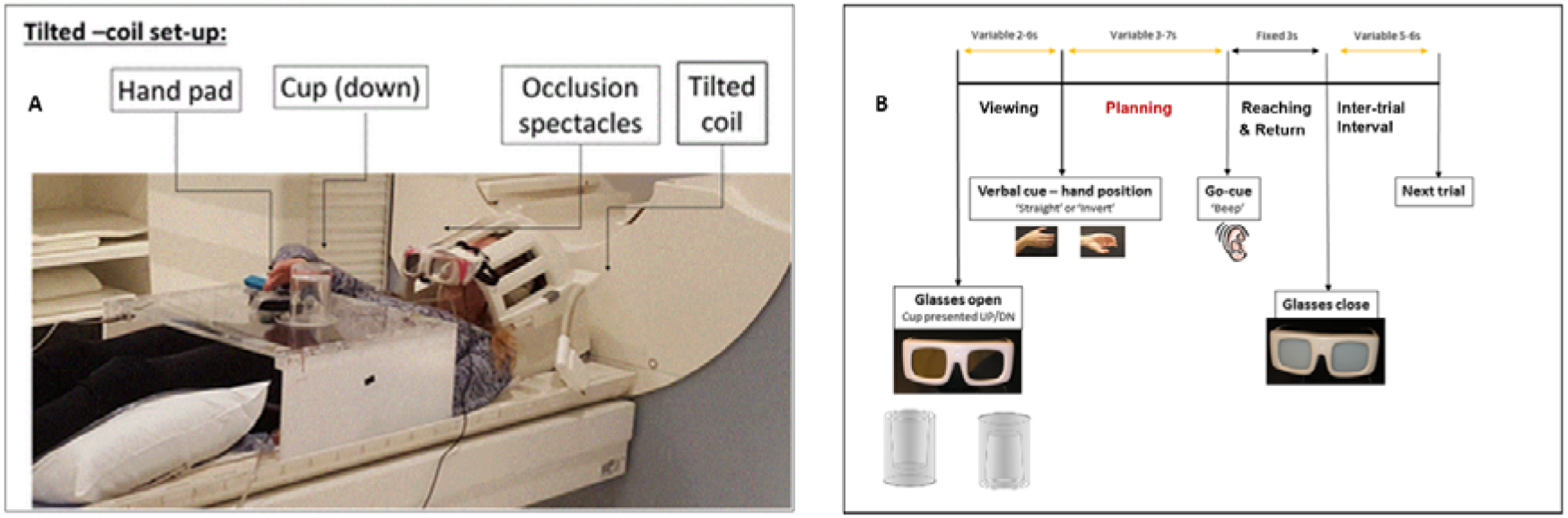
fMRI set-up and timings. (A) Experimental set-up Example of set up from one participant (video of example trial in Supplemental Material). The participant laid supine with their head on a 12-channel tilted coil (external tilt angle provided by 15° wedge, see Supplemental Material; with another 10° tilt provided by padding in the head coil). They wore PLATO occlusion spectacles which were positioned at the edge of the coil here for ease of visualisation. The participant is pictured lifting their hand from the home key (black button) to pick up the cup that is positioned upside down and turn it upright, on the cup holder. (B) Timings The viewing phase started with PLATO glasses turning transparent. After a variable delay, a verbal instruction lasting 0.5s followed which said ‘straight’ or ‘invert’ indicated the start of the Planning phase, during which participants maintained their hand on the home key until they heard a go-cue (a ‘beep’, also lasting 0.5s), which followed after a variable delay from the verbal instruction. At this point (‘Reaching’ time) participants lifted their hand from the home key (reaction time) to reach and turn the cup before returning to the home key. They were instructed to complete the action within a fixed time of 3 seconds, indicated by the glasses becoming opaque. Modelling of the GLM based on these timings is detailed in the text. The imaging results presented in this study relate to the planning phase, highlighted in red.

Participants performed actions with their right hand and had the upper arm immobilised using a wedge-shaped elbow foam pad positioned against their side and the side of the scanner bore, in front of the Perspex table which was positioned above their lap, and was secured with pegs that were fitted in the side of the scanner bed (Figure 1). The pad and Perspex table constrained participants’ arm movement to rotate around the elbow, and wrist. A ‘home’ key and cup (target object) were positioned on the Perspex table. The cup was positioned on a cup-holder that formed a dent on the Perspex table. This and the home key button were fitted with sensors allowing the measurement of times at which the home key was pressed, or the cup was lifted from (and reposition onto) the cup-holder.

A custom-made handle-less transparent cylindrical cup, measuring 10.5cm in height and 7.8cm in diameter and shaped to be perceived as upright or down was positioned on the cup holder. Participants were instructed to rest their hand on the home key button all the time except when they were due to perform an action. This allowed the measurement of the action initiation and its ending, when the hand was lifted from its resting position on the home key, to its return after having turned the cup as instructed. The cup holder was positioned at an average distance of 50cm from the participants’ eyes, adjusted to match each participant’s arm length such that all movements were comfortable (Culham et al. 2004, Gallivan et al. 2011). The cup subtended a vertical visual angle averaging 10^0^ in front of participants at a point corresponding to each participant’s sagittal midline. The home key was positioned an average of 20cm to the right side of the cup.

The timely appearance of the cup was controlled by liquid-crystal MRI-compatible, ‘PLATO’ occlusion spectacles (Translucent Technologies, Toronto, Ontario, Canada), which participants wore throughout the experiment. These allowed the timely initiation and end of each trial, and obstructed participants’ vision between trials. Experimental conditions, timings and recording of movement-related responses were controlled using a personal computer running Presentation 15.0 (Neurobehavioral Systems, San Francisco, CA).

Of note, our choice of using a handle-less cup in this experiment was to remove a confound that has led to previous debates as to whether affordance effects relate to visuo-spatial attention (cf. the ‘Simon’ effect’; Simon, 1969) or whether it constitutes the elicitation of motor representations (Wilf et al. 2013, Cho and Proctor 2013). In our task the congruency effect was not specified by the handle of a cup, but rather by its position being upright or down, which would habitually elicit a supinated or pronated grasp, respectively (Herbort and Butz 2011, Rounis et al. 2017, Pizzamiglio et al. 2020). Previous studies have demonstrated hand-object compatibility effects differ according to whether the object location is centred (Cho and Proctor 2013, Bub et al. 2018). There is literature to explain these behavioural effects in terms of differences between ‘motor’ and ‘orienting’ attention, the former being elicited when single objects are presented at the centre of vision removing confounds of oculomotor and visuo-spatial responses (Rushworth et al. 2001, Rounis et al. 2007). Motor attention involves dorsal visuomotor networks centred in the anterior parietal region and is left-lateralised with deficits leading to ideomotor apraxia (Rushworth et al. 1997). Based on this, we conjectured that affordance effects obtained from this object would not be attributable to an orienting process because the object and responses in our task were centrally located (Rounis et al. 2007, Bub et al. 2018).

### 2.3 Experimental time course and procedures

In this task, participants had to grasp the cup with their right hand and turn it either from an upright orientation to upside-down or vice versa. An event-related design averaging 8-16 seconds per trial was used to isolate visuomotor response for planning from motor execution responses, as has been done in other object-in-the-scanner experiments (Gallivan et al. 2011, 2013). Each trial was preceded by a variable period (with a variable inter-trial interval of 5~6s) in which participants had the spectacles switched off (opaque), and their right hand resting on the home key. This time allowed an experimenter, who was with the participant in the scanning room, to position the cup on the cup holder either in an upright or upside-down orientation according to a random order of conditions (determined to ensure equal repetitions of each trial type). During a trial, the experimenter was never visible to the participant when the glasses were open. The experimenter monitored performance in each trial and recorded any errors. Each trial condition was provided to the experimenter from instructions presented on a screen that was visible to them from the control room. These were not visible to the participants lying in the scanner bed.

Each trial began with the liquid spectacles turning transparent (open), allowing the participant to visualise the cup either in its upright or upside-down position (the ‘Viewing’ phase). After a random time-interval of 2~6s a verbal cue instructed the participant to either grasp the cup with a pronated or a supinated grasp. The verbal cue lasted 0.5s and consisted of the word ‘invert’ (for pronated grasp) or ‘straight’ (for supinated grasp). This verbal cue corresponded to the onset of the ‘Planning’ phase. During both these intervals, participants continued to rest their hand on the home key. Following a further variable duration of 3~7s, a beep signal (0.5s duration) was delivered. This corresponded to a ‘go cue’ indicating that participants had to execute the movement instructed in the ‘Planning’ phase. At the go-cue, participants lifted their right hand from the home key as quickly as possible, to reach and grasp the cup either with a ‘straight’ or an ‘inverted’ grasp (in the manner instructed by the verbal cue at the planning phase) and executed the action before returning to the home key, within a fixed interval of 3s, denoted when the translucent spectacles turned opaque. When participants heard the go cue, they executed the cup manipulation task instructed by the verbal cue presented during the ‘Planning’ phase. If they heard ‘straight’, participants grasped the cup using a ‘thumb-up’, supinated, wrist posture and turned it with a pronation, leading to an uncomfortable ‘thumb down’ position. If they heard ‘invert’ during the ‘Planning’ phase, participants grasped the cup using a ‘thumb-down’ pronated wrist posture and turned it with a supination to a comfortable ‘thumb up’ end state. The execution phase ended with the spectacles becoming opaque (closed) after a fixed duration of 3 seconds, before a further variable inter-trial interval of 5-6 seconds followed, during which the cup was repositioned by the Experimenter, according to the next trial’s condition.

The variable durations for each phase mentioned above (namely, the intertrial interval, ‘Viewing’, and ‘Planning’ phases) were drawn from a geometric distribution (p=0.2) in steps of 0.5s. The reason for introducing a variable time between each of these intervals was to make the auditory instruction unexpected, based on behavioural pilots.

Participants completed 10 runs of 24 trials each (4 conditions × 6 trials per condition) in one fMRI session (total of 240 trials), lasting 45-60min. The order of the trials was randomised across each run and each participant, balanced across conditions. Prior to the beginning of the scanning session, participants trained on the task for 15min outside the scanner until they were error-free and able to complete the movement execution within the 3s between the go-cue and closure of the PLATO spectacles.

### 2.4 Experimental conditions

There were four experimental conditions, in a 2 × 2 experimental design, based on the cup, and hand orientations, instructed by the task. The combination of the cup and initial hand orientation, which were provided in the ‘Planning’ phase, determined ‘affordance’ effects. The initial cup orientation was either upright or upside-down. A verbal instruction specified how participants should orient their hand grasp from the resting position on the pad after the beep. This instruction was either to orient their hand ‘straight’ or ‘invert[ed]’. This instruction determined the hand posture to adopt when grasping the cup <u>at the start of the turn</u>. Of note the hand orientation adopted at the start of the turn also determined whether the end posture was comfortable or not. The use of different verbal instructions for the hand posture (‘straight’, meaning that participants had to grasp the cup with a supinated hand posture; versus ‘invert’, meaning that they had to grasp it with a pronated hand posture) was used to prevent confounds caused by a visual instruction, such as a marker on the object, which had been used in previous versions of this experiment, published elsewhere (Rounis et al. 2017). Indeed, a visual marker to indicate the starting hand posture to use on the cup would sometimes be in conflict with the object orientation and confound any hand-cup congruency activations in an fMRI experiment. As a result of this design, the end state comfort effect at the Planning phase was influenced by the different verbal instruction cues. Although these activations are reported in the results section (in terms of a main effect of hand orientation - ‘straight’ versus ‘invert’), this was not an effect of interest in our imaging results. The effects of end state comfort have been described elsewhere (Zimmermann et al. 2013).

The combination of cup orientation and task instruction led to a congruency between the cup and hand orientation in two out of four conditions (Figure 2), which were our conditions of interest. These were the conditions when the cup was upright, and the hand instruction was ‘straight’ or when the cup orientation was <u>down</u> and the hand instruction was ‘invert’. Conversely, the two remaining conditions involved a hand orientation, specified by the verbal cue, that was incongruent with the cup orientation, such that participants grasped the closed end of the cup. These included the conditions when the verbal cue was ‘straight’ and the cup orientation was down; or else when the verbal instruction was ‘invert’ and the cup orientation was upright. In both cases, the action performed after the go-cue was to turn the cup from one orientation to the other. Figure 1 depicts the experimental set-up, and timings. The experimental task conditions are shown in Figure 2. A representative video of a task condition and images of our set up are further provided in the Supplemental Material.

**Figure 2:**
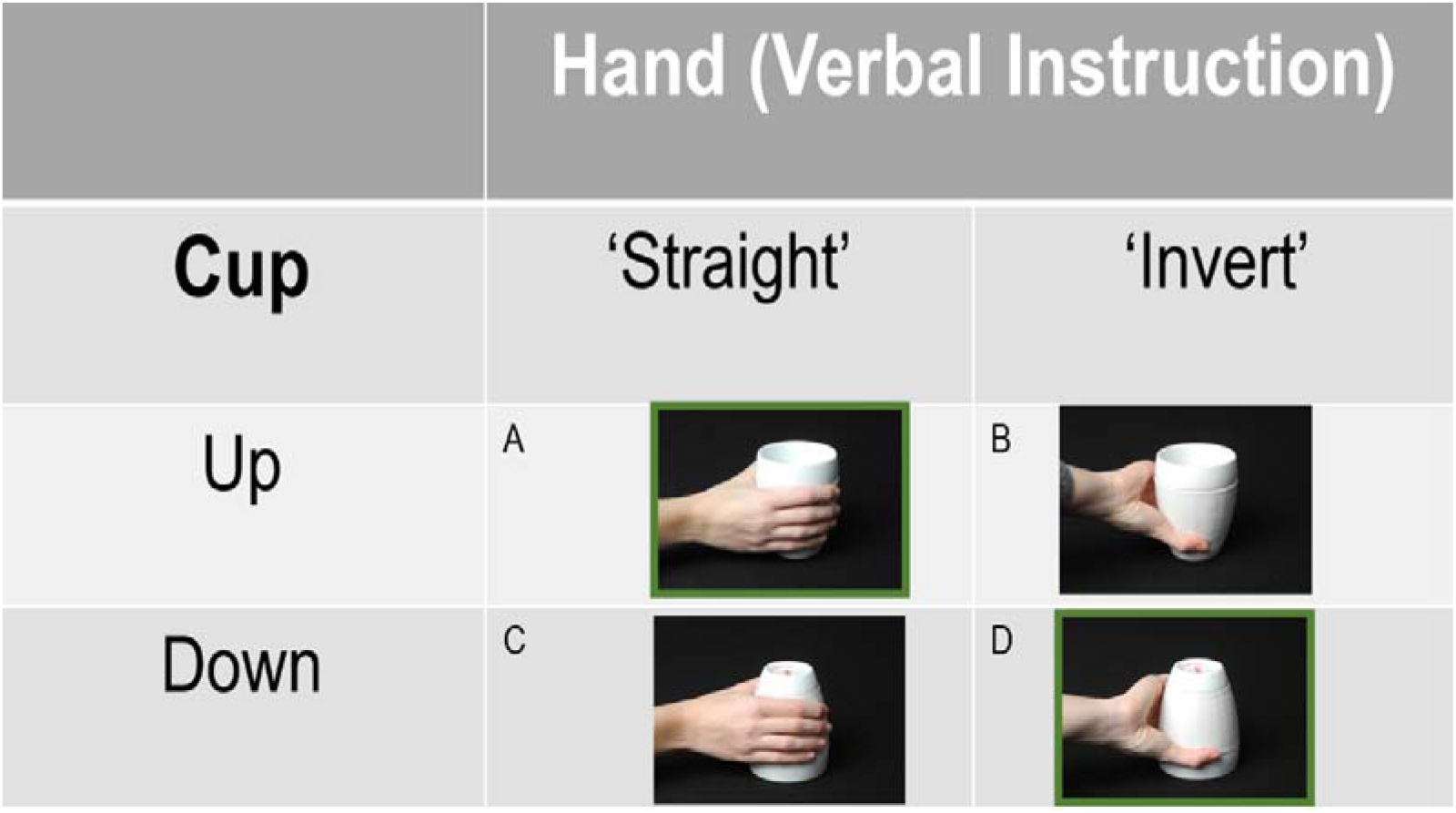
Factorial Design. This was a 2*2 factorial design: the main factors of hand and cup orientation led to an interaction of ‘affordance’ (when both were congruent, condition ‘A’ being when the cup is upright and the hand approaching it is ‘straight’ and condition ‘D’ being when the cup is upside down and the hand approaching it is ‘invert(ed)’ in this case), highlighted with a green square. The remaining, non-afforded, conditions were ‘B’ when the cup was up and the hand approaching it was inverted (note this led to a comfortable end posture after a turn) and ‘C’ when the cup was oriented down and the hand orientation at the start was straight. Of note the actual cup used in this task was purpose built with Perspex and cylindrical in shape so that its width at the top and bottom was the same, as in Figure 1.

### 2.5 Behavioural analysis

The behavioural responses relevant to the task, which are reported below corresponded to the time interval recorded between the go cue and the hand releasing the home key (the ‘reaction time’, RT), measured for each trial. This time interval is felt to represent movement planning (Wong et al 2015) and corresponds to the time at which compatibility effects in response to handled objects have been observed previously (Bub and Masson, 2010, 2011, Rounis et al. 2017, Pizzamiglio et al. 2020). RTs for each participant in each condition were entered as our dependent variable in the behavioural analyses. The remaining times (namely the time to reach the and manipulate the object, and return to the home key, i.e. from cup lifting to be turned, to cup being re-positioned back on the cup holder to hand return to the home key) were not further analysed behaviourally. However, these timings were taken into account and modelled separately from the initial parts of the movement, in the General linear model (GLM) imaging analysis.

Error trials were recorded by the experimenter who documented if the object manipulation was correctly performed in each trial, during the experiment. These included technical errors and behavioural errors (wrong grasp, action too slow, hesitation, hand posture adjusted during reaching, etc.). In addition, trials in which RTs were either above or below 2.5 SD of mean RT, or where participants took longer than 3 seconds to complete the cup manipulation, were excluded as errors. Error trials were excluded from behavioural analyses and modelled separately in the GLM imaging analysis.

A repeated measures ANOVA using RTs to investigate the effects of cup and hand congruency was implemented using IBM SPSS Statistics 25 for Windows software (SPSS Inc., Chicago, IL). As mentioned above, the effect of ‘affordance’ is equivalent to an interaction effect between Hand and Cup orientations. The type I error rate was set at p<0.05 for the analyses reported here. Greenhouse–Geisser correction for degrees of freedom was used when the assumption of sphericity was not met.

### 2.6 Image acquisition

MRI data were acquired on a Siemens 3T Trio MRI scanner at the University of Oxford Centre for Clinical Magnetic Resonance Research (OCMR). For purposes of co-registration with functional data, structural T1-weighted MRI images were acquired using the MP-RAGE sequence (repetition time, 2040ms; echo time 4.7ms; field of view 174× 192mm^2^; 192 slices; voxel size, 1×1×1mm^3^). Functional images were acquired using an echo planar imaging (EPI) sequence (repetition time, 2230ms; echo time, 30ms; flip angle, 87 degrees; isotropic voxels of 3mm, no slice gap; field of view, 192×192mm^2^; 37 slices; voxel size, 3×3×3mm^3^).

### 2.7 Imaging data pre-processing and analyses

Pre-processing: Functional imaging data were pre-processed and analysed using SPM12 (http://www.fil.ion.ucl.ac.uk/spm). The first three volumes for each session were discarded to allow for MRI signal equilibration. The image time series were spatially realigned using rigid body transformation and a sinc-interpolation algorithm (Friston et al. 1995). The time series for each voxel was temporally realigned to the first slice of each image volume.

The anatomical image was co-registered with the mean functional image, and then segmented. Deformation fields were obtained from the segmentation step, which were used to normalise the functional images to the MNI standard space. Spatial smoothing was applied to the normalised functional images with an 8-mm FWHM (full-width at half maximum) Gaussian kernel.

General Linear Model: For each participant, the fMRI time series were concatenated from 10 runs for GLM analysis (Friston et al. 1996). Single subject models consisted of regressors separately describing the ‘Viewing’ phase (glasses opening, leading to visualisation of the cup), ‘Planning’ (indexed by verbal instruction cue), ‘go-cue’ (corresponding to the auditory beep indicating action initiation), ‘Movement completion’ phase (from reaching to turn the cup to the return of the hand to the home key), ‘PLATO closure’ phases as well as the errors for all conditions. The ‘Planning’ phase was split into distinct parametric modulators for grasping movements according to cup orientation (upright or down), hand orientation (‘straight’ or ‘inverted’) and affordance (‘afforded’ being when the hand and cup orientation were congruent, ‘not afforded’ when they were not). The ‘Viewing’ phase regressor was time locked to the opening of the PLATO glasses, with a duration of zero. Each of the three ‘Planning phase’ parametric modulators were time-locked to the onset of the verbal cue (‘straight’ or ‘invert’), with a duration of zero. We conjectured that the neural correlates of cup and hand congruency or ‘affordance’ effects would occur during the ‘Planning phase’ and that these would correspond to the RT changes identified behaviourally (Wong et al. 2015, Rounis et al. 2017, Pizzamiglio et al. 2020). Moreover, affordance effects identified at the planning phase would not be confounded by movement related activity changes during motor execution. The ‘go-cue’ was modelled as a separate single regressor, with a duration of zero. The ‘movement completion phase’ was time-locked to the onset of the ‘go-cue’ and duration from hand lift-off to return to the home key after turning the cup in each trial. There was a regressor time locked to the closure of the PLATO glasses; and a duration of zero. The final regressor was for error trials, with the onset being the opening of the glasses and duration being the closing of the PLATO glasses for each error trial.

These regressors were convolved with a canonical haemodynamic response function (HRF) without derivative terms. Head motion was accounted for by adding the six head motion parameters as additional ‘nuisance’ regressors (Friston et al. 1996). Regressors that modelled the onset and duration for each run were added to account for brain activity differences across runs. Slow signal drifts were removed by using a 1/128Hz high-pass filter. Serial correlations were accounted for with an autoregressive AR (1) model.

In order to obtain the activity maps for the ‘Planning’ phase, the subject-level contrast images for each phase were subjected to a group-level random effects analysis. One-sample t-tests were used to compare between conditions of interest. We assessed the effects of the Hand, Cup and Affordance by using subject-level contrast images for the parametric modulators in group-level one-sample t-tests. We applied cluster-wise family wise error (FWE) corrected for multiple comparisons at p < 0.05, with a height cluster-forming threshold of p<0.001 across the whole brain.

Changes in connectivity within the network engaged in this task were assessed using ‘Psycho-physiological Interactions’ (PPI), a method first described by Friston et al (1997). The PPI analysis explains responses in one cortical area in terms of an interaction between activity in another cortical area (index area) and the influence of an experimental condition. We used this to test the hypothesis that congruency between the hand and cup orientations specified during the task instruction modulated connectivity between the left PMd, involved in motor planning, with other areas involved in the Planning phase of this cup manipulation task. This hypothesis is based on previous literature which reports this area to be involved in motor planning for object use (Grafton et al. 1998, Grezes et al. 2003, Gallivan et al. 2011 and 2013) and more specifically in representing affordances (Grezes et al. 1998, Cisek and Kalaska 2005, 2007, 2010). Three variables were created for this PPI analysis in a generalized linear model (GLM): a physiological variable for the BOLD signal in the seed region, a psychological variable corresponding to the parametric modulator for the congruency effect at the planning phase, and a psycho-physiological interaction variable. The seed was selected based on the specific effects of congruency in that phase and a-priori hypothesis for a role of dorsal premotor areas in action selection (Grafton et al. 1998, Cisek and Kalaska 2005, Arbib et al. 2000). We wanted to investigate changes in connectivity with left PMd underlying the congruency effects. For each participant, we located the peak voxel within the cluster identified by the group-level ‘congruency’ contrast for the ‘Planning’ phase and built a 6mm sphere VOI centred at the peak voxel. We extracted BOLD signal from each VOI, adjusted for the effects of the Hand, Cup and Congruency at the planning phase. In order to derive brain interactions at the neuronal level, the BOLD signal was deconvolved through haemodynamic function onto the neural level before creating the interaction variable. These three PPI variables were fed into a GLM analysis, together with six head motion estimates as variables of no interest. Subject-level contrast images for the interaction variable were entered in group-level one-sample t tests.

The anatomical localization for significant regions was identified based on the SPM anatomy toolbox (Eikhoff et al. 2005), supplemented by the multi-modal parcellation of human cerebral cortex provided by the Human Connectome Project (HCP) (Andreas, 2016; Glasser et al., 2016) and direct anatomical interpretation of our results based on Petrides’ ‘Atlas of the Morphology of the Human Cerebral Cortex on the MNI Brain’ (2018). The Figures were created using the Brain Net viewer (https://www.nitrc.org/projects/bnv/, Xia et al. 2013).

## 3. Results

### 3.1 Behavioural results

Error trials including behavioural errors (1.85%) and technical errors (2.35%) were excluded from RT analysis. Outlier RTs were removed based on 2.5 standard deviations from the mean value for each condition and trials which were completed beyond 3 seconds for each subject (2.52% excluded). Hence the total number of trials (403) excluded were 6.72% of all trials. Error Trials were not analysed any further.

A repeated measure ANOVA for the RT data (Figure 3) revealed a significant main effect of hand posture (F(1,24)=46.5, p=4.7E-07, partial eta^2^=0.66, MSE 228.733), with ‘inverted’ grasp being initiated with shorter RTs than ‘straight’ grasps (501.48ms vs. 522.11ms), no main effect of cup orientation (F(1,24)=0.124, p=0.728, partial eta^2^=0.005, MSE = 295.56); and a significant interaction between the two (F(1,24) = 7.551, p=0.011, partial eta^2^=0.24, MSE= 367.21), with shorter RTs for the conditions in which hand and cup orientations were congruent (‘afforded’ trials – Figure 2) than ones in which they were not (506.53ms vs. 517.01ms).

**Figure 3:**
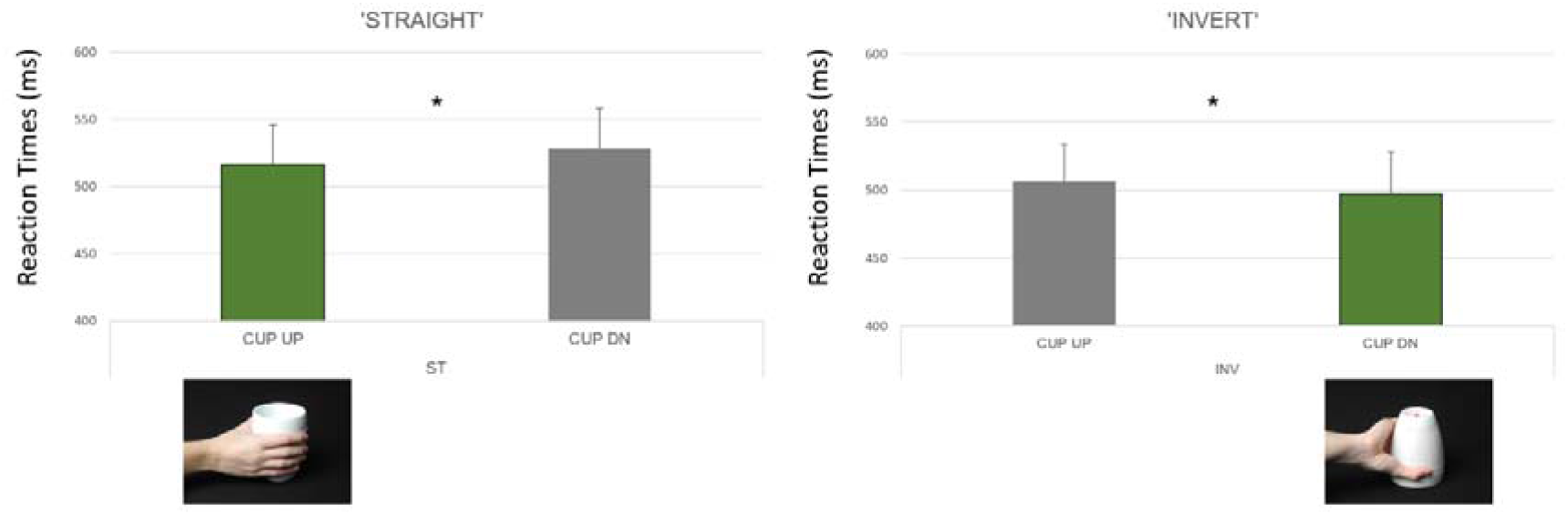
Behavioural results. Reaction Times Figure depicting the behavioural results. The reaction times (RTs) represented the time at which participants lifted their hand off the home key to initiate the action. The left panel reports RTs for actions that were initiated with a ‘straight’ (supinated) hand posture when reaching to turn the cup. The right panel reports RTs for actions that were initiated with an ‘inverted’ (pronated) hand orientation. In both cases, we identified effects of congruency between the hand and cup orientations at this time, indicating that RTs were shorter for trials in which the hand and cup orientations were congruent than for ones in which they were not (*p=0.011 – effect of ‘affordance’); moreover, they were shorter when planning actions starting with an inverted grasp and ending comfortably, compared to ones which started with a ‘straight’ hand orientation (***p=4.7E-07).

### 3.2 Imaging Results

A random-effects analysis investigating effects of our task conditions at the group-level was performed. The overall activations at the Planning phase, relative to the implicit baseline of inter-trial intervals, are reported in Figure 4 (left) and in Table 1. The results reported here were whole-brain corrected at FWE p<0.05, cluster-wise.

**Figure 4:**
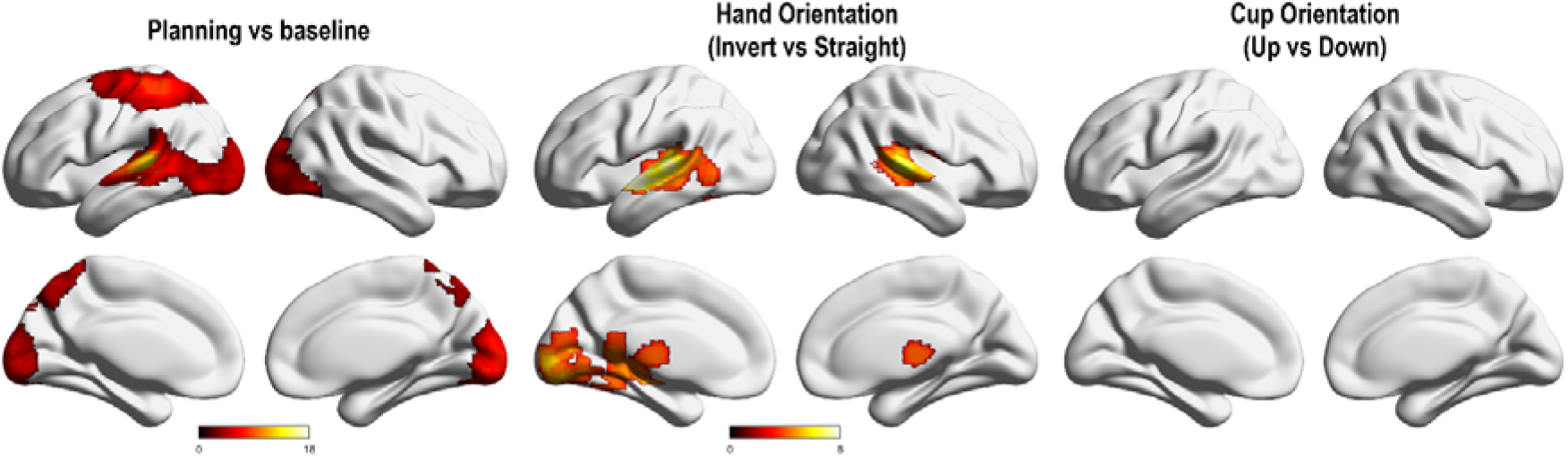
Imaging results at the planning phase. This figure depicts activation maps in Planning versus baseline (the activation reported in Table 1) on the left, effect of hand orientation in the middle– depicting the conditions when the hand instruction was straight (leading to uncomfortable end state) versus the ones in which it was ‘invert’ (leading to comfortable end state), and effects of cup orientation on the right, with no significant activation for that condition. The results are shown at FWE<0.05 whole brain, cluster-wise correction. The activation maps have been overlayed on a rendered structural T1 MRI map in MNI space from BrainNet viewer (https://www.nitrc.org/projects/bnv/), depicting activations in lateral (top panels) and medial (bottom panels) aspects of the left and right hemispheres, respectively. The colour bar indicates T values for activations in the areas of interest.

**Table 1:** Brain regions associated with increased activity during the Planning Phase (at time of Verbal Cue Instruction) (cluster-wise pFWE<0.05)

| Anatomical Region | Hemisp<br>here | X | Y | Z | T<br>value | Voxel<br>count | cluster-level p<br>FWE-correcte<br>d |
| --- | --- | --- | --- | --- | --- | --- | --- |
| Middle Temporal Gyrus | Left | -60 | -34 | 8 | 12.40 | 1514 | <0.001 |
| Superior Temporal Gyrus | Left | -57 | -16 | 5 | 8.80 |  |  |
|  |  | -60 | 2 | -7 | 4.11 |  |  |
| Cuneus (V2, V3d) | Left | -3 | -97 | 14 | 7.29 |  |  |
| Calcarine Gyrus (V1, V2) | Left | 0 | -88 | 2 | 6.22 |  |  |
|  | Right | 6 | -97 | 5 | 7.41 |  |  |
|  |  | 3 | -88 | -7 | 7.28 |  |  |
| Inferior Occipital Gyrus<br>(Lateral occipito-temporal<br>complex) | Left | -51 | -73 | -4 | 6.33 |  |  |
|  |  | -33 | -91 | -10 | 5.88 |  |  |
|  | Right | 36 | -82 | -13 | 6.21 |  |  |
|  |  | 30 | -91 | -10 | 6.64 |  |  |
| Inferior<br>Occipitotemporal/Fusiform<br>Gyrus | Right | 48 | -70 | -13 | 5.10 |  |  |
| Middle Occipital Gyrus<br>(V3d/V3A) | Left | -21 | -97 | 11 | 5.51 |  |  |
|  | Right | 30 | -88 | 20 | 3.78 |  |  |
|  | Right | 36 | -91 | -1 | 6.37 |  |  |
| Superior Occipital Gyrus<br>(V3d/V3A) | Left | -12 | -97 | 14 | 6.25 |  |  |
|  | Right | 24 | -94 | 11 | 5.65 |  |  |
| Cerebellum | Right | 24 | -73 | -19 |  |  |  |
| Superior Parietal Lobe<br>(Area 2, 5L, 7PC) | Left | -33 | -43 | 59 | 8.37 | 921 | <0.001 |
|  |  | -24 | -55 | 65 | 5.89 |  |  |
|  |  | -15 | -73 | 53 | 4.64 |  |  |
| Postcentral Gyrus (Area 4p, 4a, 3b) - Primary motor area (M1) | Left | -36 | -34 | 59 | 8.28 |  |  |
| Precentral gyrus, dorsal premotor cortex | Left | -33 | -19 | 68 | 6.64 |  |  |
|  |  | -36 | -7 | 65 | 6.25 |  |  |
|  |  | -24 | -7 | 65 | 4.91 |  |  |
| Precuneus/ Superior Parietal Lobule (7P/7A) | Left | -3 | -52 | 68 | 6.97 |  |  |
|  |  | -6 | -79 | 47 | 5.93 |  |  |
|  |  | -3 | -64 | 59 | 5.45 |  |  |
|  | Right | 9 | -70 | 59 | 4.55 |  |  |
|  |  | 9 | -76 | 53 | 4.22 |  |  |

A wide network of areas was activated, predominantly within the precentral, postcentral gyri, superior parietal lobule, intraparietal sulcus of the left hemisphere but also including bilateral activations in the occipital and temporal areas. The left superior temporal activation included auditory and visual subdivisions (notably BA22) and adjacent left middle temporal gyrus (Figure 4, Table 1).

The main effect of hand orientation (which was represented by the initial hand orientation being ‘inverted’ for a comfortable end-state, versus ‘straight’ for an uncomfortable one) activated bilateral superior temporal gyri (including auditory areas, corresponding to the auditory cue instruction, and visual subdivisions BA22), occipital cortices including inferotemporal and lateral occipito-temporal areas and thalamus (Supplementary Table 1, Figure 4, middle panel). Activity in these areas was greater when turning a cup with an inverted (pronated) grasp, to end in a comfortable, supinated, hand posture, compared to turning it with a straight (supinated) grasp to end in an uncomfortable, pronated posture.

There were no significant activations identified for the main effect of cup orientation at the Planning phase (Figure 4, right panel).

The interaction between the cup and hand orientations, namely the effect of ‘affordance’ in the ‘Planning’ phase, revealed significant activations in the left and right dorsal premotor cortices (L PMd main cluster x=-24, y=-7, z=59, T=4.40, cluster size 89 voxels, pFWE=0.015; R PMd main cluster x=21, y=2, z=56, T=5.20, cluster size 88 voxels, pFWE=0.015). The sign of this congruency effect indicated greater activation for trials that were ‘not’ afforded, i.e. where hand and cup orientations were incongruent (Table 2, Figure 5).

**Table 2:**
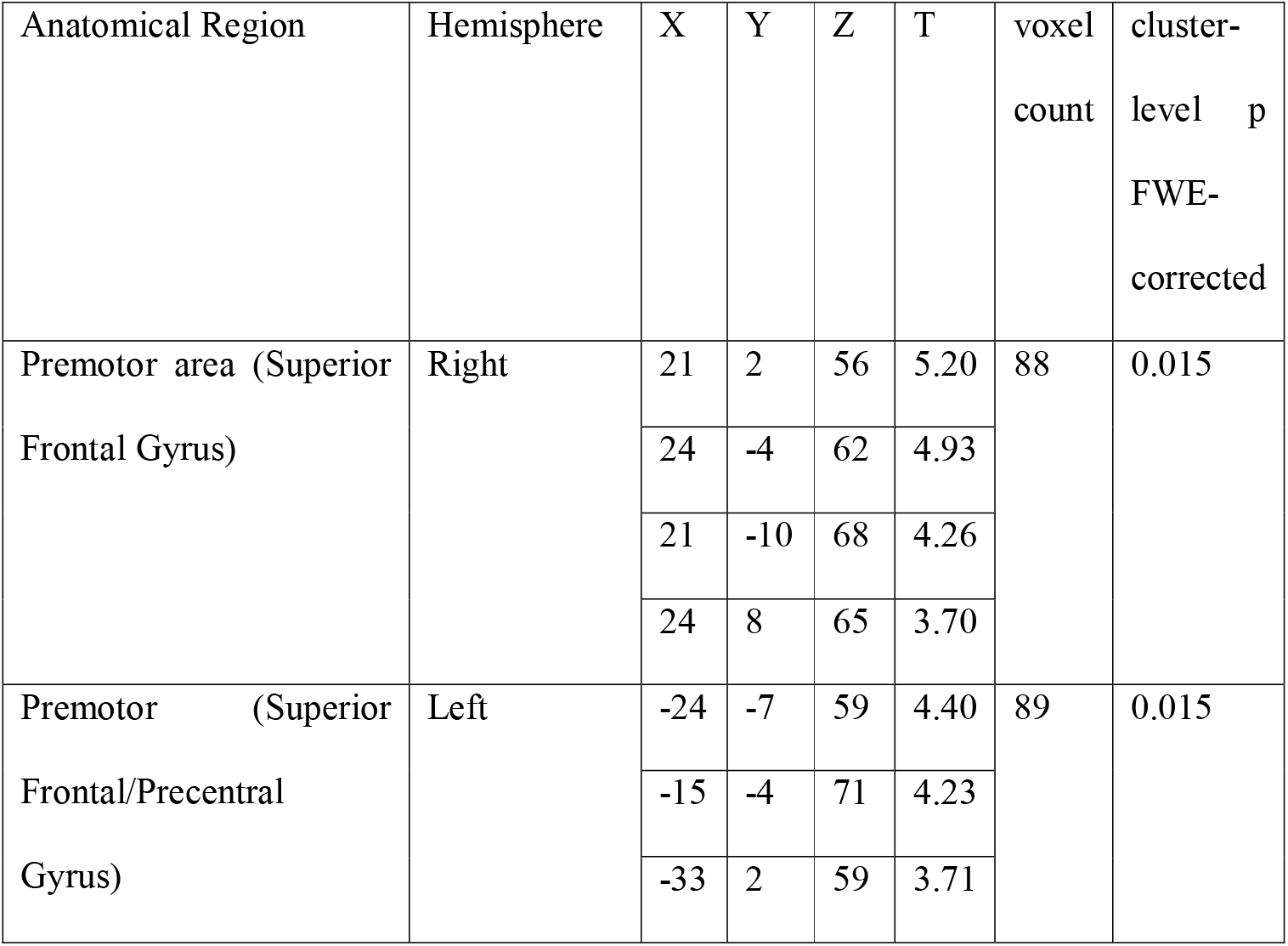
The effects of congruency (incongruent>congruent) during the Planning phase (cluster-wise, pFWE<0.05)

| Anatomical Region | Hemisphere | X | Y | Z | T | voxel count | cluster-level p FWE-corrected |
| --- | --- | --- | --- | --- | --- | --- | --- |
| Premotor area (Superior Frontal Gyrus) | Right | 21 | 2 | 56 | 5.20 | 88 | 0.015 |
|  |  | 24 | -4 | 62 | 4.93 |  |  |
|  |  | 21 | -10 | 68 | 4.26 |  |  |
|  |  | 24 | 8 | 65 | 3.70 |  |  |
| Premotor (Superior Frontal/Precentral Gyrus) | Left | -24 | -7 | 59 | 4.40 | 89 | 0.015 |
|  |  | -15 | -4 | 71 | 4.23 |  |  |
|  |  | -33 | 2 | 59 | 3.71 |  |  |

**Figure 5:**
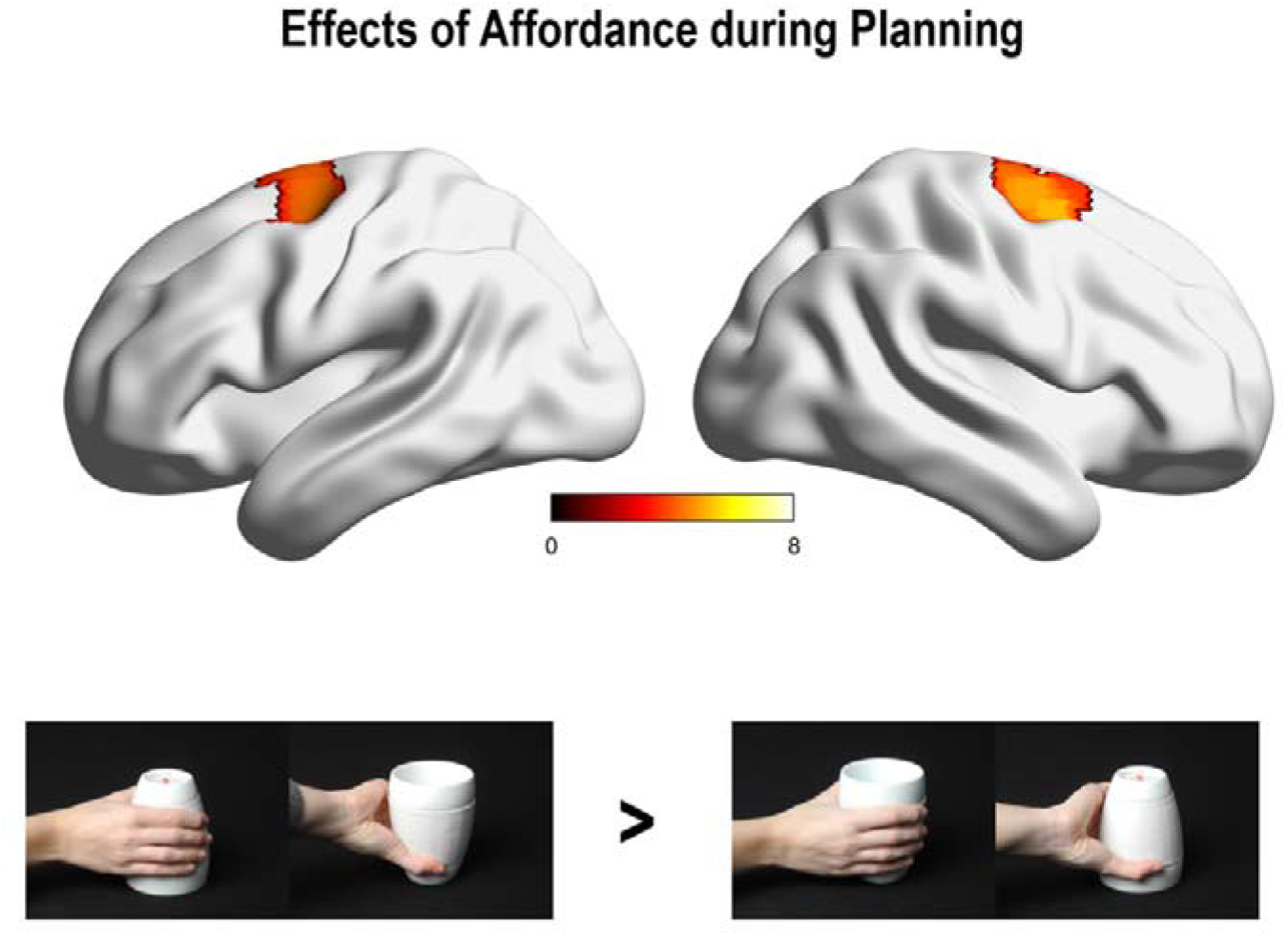
Effects of Hand-Cup Congruency on task related activity. Activation map for the effect of affordance at the Planning phase. The results are shown at pFWE<0.05 cluster-wise correction. There was significantly increased activity in the left and right dorsal premotor (PMd) cortices in conditions in which the hand and cup orientation were incongruent during the Planning phase.

We then applied PPI analyses to test the hypothesis that congruency between the hand and cup orientations specified during the task instruction modulated connectivity between the PMd areas identified as mediating the ‘affordance effect’ in this and previous studies (Grezes et al. 2003, Cisek and Kalaska 2005, 2007, 2010) and other areas involved in the Planning object manipulation within dorsal and ventral stream (Grafton et al. 1998, Grezes et al. 2003, Gallivan et al. 2011 and 2013, Sakreida et al. 2016).

The left PMd (x=-24, y=-7, z=59) involved during movement planning was chosen as the seed area for our PPI analysis, looking for changes in coupling between this area and areas of the dorsal and ventral visuomotor networks based on hand-object congruency, during the planning phase. This PPI revealed one area in which coupling was significantly increased in conditions that were incongruent within the left lateral occipito-temporal cortex (x=-30 y=-85 z=-10, T=5.29, and x=-39, y=-76, z= −4, T=4.9, cluster of 71 voxels, pFWE=0.028). Coupling between the left PMd and LOTC area was increased when the hand and cup orientations were incongruent (Figure 6). Of note, a PPI investigating affordance-related connectivity changes with the right PMd (x=21, y=2, z=56), revealed no significant results.

**Figure 6:**
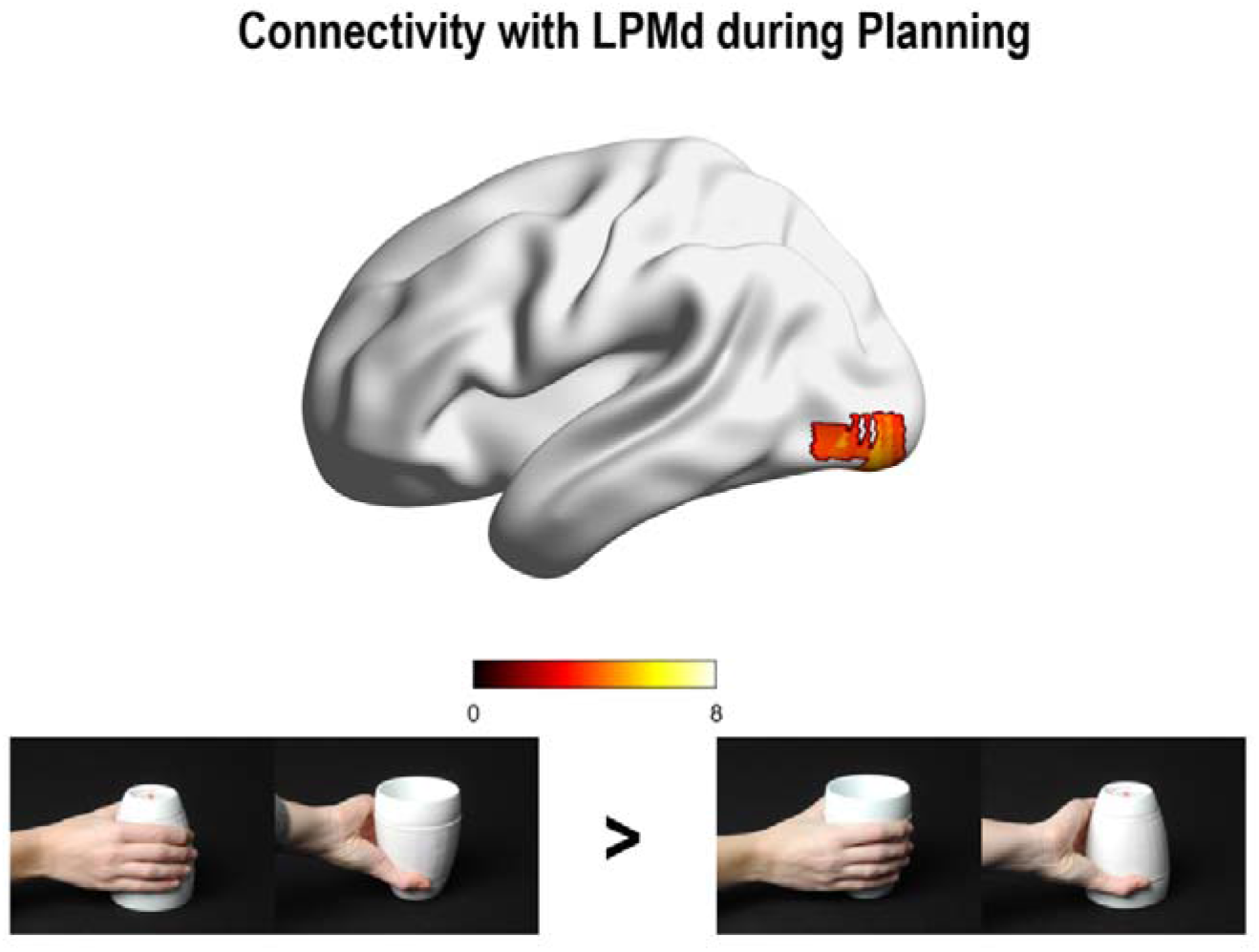
LPMd PPI Results. Activation map identifying areas of increased connectivity with the left PMd modulated by affordances in the Planning phase. The areas included formed part of the left inferotemporal and lateral occipital areas forming the lateral occipito-temporal cortex (LOTC). As in the previous figure, these activation maps have been overlayed on a rendered structural T1 MRI map in MNI space from BrainNet viewer and the colour bar indicates T values for activations in the areas of interest.

## 4. Discussion

In this study, we investigated the influence of cup orientation on goal-directed actions when planning to turn it. To our knowledge this is the first study pitting congruent versus incongruent hand-object interactions during real object manipulation in a functional imaging environment. Participants performed a delayed-movement task in which they reached and turned a cup either when it was oriented upright or upside down. They were instructed to use a hand orientation to turn the cup that was either congruent or incongruent with the object orientation. As in previous studies (Rounis et al. 2017, Pizzamiglio et al. 2020), we identified a behavioural effect of congruency (an ‘affordance’ effect). Movements where the hand and cup orientations were matched were initiated faster than ones in which they were not.

Planning to turn a cup in this task affected activity in areas involved in object manipulation (Goodale and Milner 1992, Rizolatti and Matelli 2003, Grezes et al. 2003, Drapati and Sirigu 2006, Mahon et al. 2007, Grol et al. 2007, Gallivan et al 2011, 2013). In addition, there was increased activity in bilateral PMd for trials that were not afforded. The left PMd’s connectivity with the left LOTC was increased in those same trial conditions. We discuss these imaging and behavioural results in the context of previous literature investigating motor control and hand posture representations and propose that they reflect processes underlying selection of hand postures for a task.

### The neural correlates of affordances on posture representations in the brain

In this study, planning object-related manipulations in which object and hand orientations were incongruent was associated with increased activity in bilateral PMd. PPI analysis investigating areas of connectivity changes relating to this congruency effect, revealed increased functional connectivity between left PMd and the left LOTC in incongruent (ie. ‘non-afforded’) task conditions.

The PMd plays an important role in action selection (Grafton et al. 1996, Grafton et al., 1998; Rumiati et al. 2004, Cisek, 2007), within the dorsal stream network (Fagg and Arbib 1998). Neurophysiological studies in non-human primates have shown that LPMd activity increases during motor preparation when planning competing reach movements (Cisek, 2006, 2007; Cisek & Kalaska, 2005). Taken together, there is evidence suggesting that PMd may be involved in triggering a competition process elicited by affordances in humans. Applying repetitive transcranial magnetic stimulation (rTMS) over this area leads to slower motor performance when the instructed response is not congruent with the visual stimulus (Praamstra et al., 1999; Rushworth et al., 2001, Glover et al. 2005 Ward et al. 2010, Makris et al. 2011).

A previous imaging study investigated the neural correlates of affordances using Tucker and Ellis’ (2001) paradigm (Grezes et al. 2003). Participants in that task had to categorise objects as either natural or man-made by making a precision grip for one category and a power grip for another, in a counterbalanced order. They identified ‘affordance’ or compatibility effects to be associated with areas within both ‘dorso-dorsal’ and ‘ventro-dorsal pathways’ (Rizzolatti and Matelli 2003). These included the anterior intraparietal area, PMd and inferior frontal cortex. The anterior intraparietal and inferior frontal areas are known to be involved in grip selection (Fagg and Arbib 1998) and may incorporate more conceptual information for object use (Drapati and Sirigu 2006, Van Polanen and Davare 2015). The left PMd is located in the ‘dorso-dorsal’ pathway for action selection and reaching; its role in affordances corroborates our results. The stimuli used for eliciting affordances in the Grezes et al. (2003) task involved 2D images, compared to a real object (a cup) in ours. Moreover, their task involved object categorisation. Previous studies have reported stronger affordance effects with real objects (Snow et al. 2011, Gomez et al. 2017) compared to 2D images of objects (Bub and Masson 2010, Squires et al. 2016, Bub et al. 2018). In a recent study, the categorisation of real objects led to the use of factors relating both conceptual and physical characteristics, whereas 2D-images were mostly categorised on the basis of conceptual characteristics alone (Holler et al. 2020). Taken together, these differences might explain differences in activation patterns identified between the Grezes et al. (2003) grasp categorisation study, and ours, which involved turning a real object with a reaching and wrist rotation movements.

In addition to enhanced activity in dorso-dorsal PMd areas, planning incongruent hand-object actions was associated with functional connectivity changes between the left PMd and ventral stream area LOTC. The inferotemporal area and adjacent inferior occipital lobe, form the ventral stream pathway representing objects (Dolan et al. 1997, Kanwisher et al. 1999, Chao et al. 1999, Mahon et al. 2007). This area has been shown to incorporate knowledge of body and hand posture for tool use (Valyear et al. 2007, Rice et al. 2007, Zimmermann et al. 2013, 2018, Bracci et al. 2010, 2013, 2018). It responds to movement invariant hand postures and to motor-element properties of objects (Bracci et al.2010, 2013, 2018, Lingnau and Downing 2015, Wurm et al. 2017) and is functionally connected with dorsal stream areas (Zimmermann et al. 2018).

Previous functional imaging studies involving object-directed actions in the scanner have also identified task-related BOLD activations within subdivisions of dorsal and ventral visual stream areas (Sakreida et al. 2016) dependent of the type of action performed (eg. grip) and properties of the object (eg. large or small). Most of these involved ‘ventro-dorsal’ fronto-parietal areas with varying degrees of dorso-dosal and ventral stream involvement (Valyear et al. 2007, Grol et al. 2007, Mahon et al. 2007, Gallivan et al. 2011, 2013, Sakreida et al., 2016, Fabbri et al. 2014 and 2016). Our results indicated functional interactions between ‘dorso-dorsal’ LPMd and ventral stream area LOCT in trials where hand posture and object orientation were incongruent, suggesting a direct integration between dorsal and ventral stream areas when preparing incongruent object manipulations (van Polanen and Davare 2015). Integrating ventral stream information during object manipulation dynamically with dorsal stream structures would suggest that affordance effects represent an influence of action semantics in tasks that do not require object categorisation or understanding, such as object manipulation (Creem and Proffitt 2001, Till et al. 2014, van Elk et al. 2014, Holler et al. 2020).

Taken together, these results might explain mechanisms underlying certain conditions that display deficits affecting both dorsal and ventral stream networks, yet do not affect understanding, such as limb apraxia (Binkofski and Buxbaum 2013, Rounis and Humphreys 2015). A recent study in which stroke patients with and without apraxia performed a similar cup manipulation task to this one, identified behavioural impairments in apraxic patients, who were unable to plan actions in which the cup and hand orientation were incongruent (Pizzamiglio et al. 2020). Based on our results, this impairment may be explained in one of two ways. One possible mechanism would be an inability to signal competing actions to select via PMd. Alternatively, patients may have a deficit in integrating alternative posture representations from LOTC in incongruent trials. Further research would be required to test these alternative hypotheses.

### Affordances or Competition between Habitual and Goal Directed Actions

This study replicated behavioural effects of hand-object compatibility, in an fMRI environment, previously observed using the same task in healthy volunteers and in stroke patients (Rounis et al. 2017, Pizzamiglio et al. 2020). Motor initiation was faster in trials in which the hand and cup orientation were congruent. Reaction times (RT) represent the time when a decision about what action to implement and how to execute it, take place (Wong et al. 2015). Several studies have reported compatibility effects at that time (Tucker and Ellis 1998, Grezes et al. 2003, Bub and Masson 2010, Rounis et al. 2017). In addition to RT effects, affordances affect kinematic measures during object directed actions in human studies (Gentilucci 2002, Rounis et al. 2018). The longer RTs and kinematic changes we and others have observed in incompatible trials might represent a competition between movement representations that are habitual (Herbort and Butz 2011; Rounis et al. 2017, 2018) compared to the ones demanded by the task. This has been demonstrated in neurophysiological studies involving non-human primates (Cisek and Kalaska 2005) as well as from neuropsychological studies involving patients with Alien Limb Syndrome (Riddoch et al. 1998, McBride et al. 2013).

Riddoch et al. (1998) studied a patient with corticobasal degeneration and alien-limb syndrome, who was asked to reach and grasp a cup using the hand that was on the same side of the table as the cup, regardless of which way its handle was oriented. The patient performed the task correctly when the cup’s handle was on the same side as the hand she was instructed to use. However, if the handle was on the opposite side, there were ‘interference’ errors: in this case, the patient was unable to inhibit the action of grasping the cup with the opposite hand, the action cued by the orientation of the cup’s handle in relation to the patient’s preferred hand. These errors were not present when she was asked to point to the object or responded to lights instead of cups, suggesting that for this patient, the object elicited an associated motor plan which was movement specific.

Taken together, our results suggest that affordances are not represented by independent neural network. Instead, they are task specific. In this task, which involved alternative wrist postures for turning a cup, affordances appeared to modulate representations of hand-object orientations during their integration with motor control (PMd) networks specifying reaching movement to the cup. Further investigations of these effects using EEG or MEG would extend findings from this neuroimaging study, to determine whether deficits identified in patients may arise from a failure in the elicitation of appropriate hand postures in LOTC, or from a failure in action selection signalled by LPMd.

## CRediT author contribution statement

**Zuo Zhang**: Formal analysis, Investigation, Methodology, Project administration, Visualization, Writing – original draft, Writing – review & editing; **Natalie Nelissen**: Methodology design, Software programming of task, Data Curation, Writing – review & editing; **Peter Zeidman**: Formal Analysis, Validation, Supervision, Analysis tools, Writing – original draft, Writing – review & editing; **Nicola Filippini**: Data acquisition, Project administration, Writing – review & editing; **Jörn Diedrichsen**: Conceptualisation, Methodology, Supervision, Writing – review & editing; **Stefania Bracci**, Software, Validation, Visualisation, Writing – review & editing; **Karl Friston**: Formal Analysis, Analysis Resources, Supervision; **Elisabeth Rounis**: Conceptualisation, Methodology, Funding Acquisition, Project Administration, Data acquisition, Supervision, Writing – Original Draft, Writing – Review & Editing.

## Acknowledgements

We would like to thank the participants who took part in the study. This study was supported by personal grants to Dr E. Rounis from the British Medical Association (Helen Lawson grant), Academy of Medical Sciences and The Oxford Charitable Trust. We would like to thank Daniel Voyce; John Prentice from the MRC Oxford Institute of Radiation Oncology; Gloria Pizzamiglio; Steven Knight and Professor R. Passingham for their help and advice with this study.

## References

Andreas H. 2016. HCP-MMP1.0 projected on MNI2009a GM (volumetric) in NIfTI format [Online]. Available: https://doi.org/10.6084/m9.figshare.3501911.v5.

Arbib MA. 1997. From visual affordances in monkey parietal cortex to hippocampo-parietal interactions underlying rat navigation. Phil Trans R Soc Lond B Biol Sci 352, 1429–1436

Arbib MA, Billard, A, Iacoboni, M & Oztop, E. 2000. Synthetic brain imaging: grasping, mirror neurons and imitation. Neural Netw, 13, 975.–997.

Balleine BW, Dickinson A. 1998. Goal-directed instrumental action: contingency and incentive learning and their cortical substrates. Neuropharmacology 37(4-5), 407–419. doi: 10.1016/s0028-3908(98)00033-1.

Bernstein N. 1967. The coordination and regulation of movement. Pergamon Press, Oxford.

Binkofski F, Buxbaum LJ. 2013. Two action systems in the human brain. Brain Lang 127:222–229.

Bracci S, Iestwaart M, Peelen MV, Cavina-Pratesi C. 2010. Dissociable neural responses to hands and non-hand body parts in human left extrastriate visual cortex. J Neurophysiol 103, 3389–97.

Bracci S, Peelen MV. 2013. Body and object effectors: the organisation of object representations in high-level visual cortex reflexts body-object interactions. J Neurosci, 33(46), 18247–58.

Bracci S, Caramazza A, Peelen MV. 2018. View-invariant representation of hand postures in the human lateral occipitotemporal cortex. Neuroimage 181, 446–52.

Brandi ML, Wohlschlager A, Sorg C, Hermsdorfer J. 2014. The Neural Correlates of Planning and Executing Actual Tool Use. J Neurosci 34:13183–13194.

Bub DN, Masson MEJ. 2010. Grasping beer mugs: on the dynamics of alignment effects induced by handled objects. J Exp Psychol Hum Percept Perform 36 2, 341–358.

Bub DN, Masson MEJ, Kumar R. 2018. Time course of motor affordances evoked by pictured objects and words. J Exp Psychol Hum Percept Perform 44 1, 53–68.

Castiello U. 1999. Mechanisms of selection for the control of hand action. Trends Cogn Sci 3(7), 857–68.

Cavina-Pratesi C, Goodale MA, Cullham JC. 2007. fMRI reveals a dissociation between grasping and perceiving the size of real 3D objects. PLoS One 2(5), 424.

Cavina-Pratesi C, Monaco S, Fattori P, Galletti C, McAdam TD, Quinlan DJ, Goodale MA, Culham JC. 2010. Functional Magnetic Resonance Imaging Reveals the Neural Substrates of Arm Transport and Grip Formation in Reach-to-Grasp Actions in Humans. J Neurosci 30:10306–10323.

Chao LL, Martin A. 2000. Representation of manipulable man-made objects in the dorsal stream. Neuroimage 12(4), 478–484. doi: 10.1006/nimg.2000.0635.

Cho D.T., Proctor R.W. 2013. Object-based correspondence effects for action-relevant and surface-property judgements with keypress resonses: evidence for a basis in spatial coding. Psychological Research 77, 618–636.

Cisek P. 2005. Neural representations of motor plans, desired trajectories, and controlled objects. Cogn Process 6(1), 15–24. doi: 10.1007/s10339-004-0046-7.

Cisek P, Kalaska JF. 2005. Neural correlates of reaching decisions in dorsal premotor cortex: specifications of multiple direction choices and final selection of action. Neuron 45(5): 801–814.

Cisek P. 2007. Cortical mechanisms for action selection: the affordance competition hypothesis. Philos Trans R Soc Lond B Biol Sci 362(1485), 1585–1589.

Cisek P, Kalaska JF. 2010. Neural mechanisms for interacting with a world full of action choices. Ann Rev Neurosci 33, 269–98.

Craighero L., Fadiga L., Rizzolatti G., Umilta C. 1999. Action for perception: a motor-visual attentional effect. J Exp Psychol Hum Percept Perform, 25(6), 1673–92.

Craighero L, Fadiga L, Umilta CA, Rizzolatti G. 1996. Evidence for visuomotor priming effect. Neuroreport 8(1), 347–9.

Creem, SH, Proffitt, DR. 2001. Grasping objects by their handles: a necessary interaction between cognition and action. J Exp Psychol Hum Percept Perform 27(1), 218–228.

Culham JC. 2004. Human brain imaging reveals a parietal area specialised for grasping. In N. Kanwisher & J. Duncan (Eds) Attention and performance XX: Functional brain – Imaging of human cognition (417–38). Oxford, UK: Oxford University Press.

Daw ND, Niv Y, Dayan P. 2005. Uncertainty-based competition between prefrontal and dorsolateral striatal systems for behavioural control. Nat Neurosci 8(12), 1704–11.

Dolan RJ, Fink GR, Rolls E, Booth M, Holmes A, Frackowiak RS, Friston KJ. 1997. How the brain learns to see objects and faces in an impoverished context. Nature 389(6651): 596–9.

Drapati E., Sirigu A. 2006. How we interact with objects: learning from brain lesions. Trends Cogn Sci. 10(6):265–270.

Eickhoff SB, Stephan KE, Mohlberg H, Grefkes C, Fink GR, Amunts K, Zilles K. 2005. A new SPM toolbox for combining probabilistic cytoarchitectonic maps and functional imaging data. NeuroImage 25, 1325–1335.

Ellis R, Tucker M. 2000. Micro-affordance: the potentiation of components of action by seen objects. Br J Psychol 91, 451–471

Fabbri S, Strnad L, Caramazza A, Lingnau A. 2014. Overlapping representations for grip type and reach direction. NeuroImage 94:138–146.

Fabbri S, Stubbs KM, Cusack R, Culham JC. 2016. Disentangling Representations of Object and Grasp Properties in the Human Brain. J Neurosci 36:7648–7662.

Fagg AH, Arbib M A. 1998. Modeling parietal-premotor interactions in primate control of grasping. Neural Netw, 11(7-8), 1277–1303.

Friston KJ, Holmes AP, Worsley KP, Poline JOB, Frith CD, Frackowiak RSJ 1996. Statistical parametric maps in functional neuroimaging: a general linear approach. Hum Brain Map 2, 189–210.

Friston KJ, Buechel C, Fink GR, Morris J, Rolls E, Dolan RJ. 1997. Psychophysiological and modulatory interactions in neuroimaging. Neuroimage 6, 218–229.

Friston KJ, Shiner T, Fitzgerald T, Galea JM, Adams R, Brown H, Dolan RJ, Moran R, Stephan KE, Bestmann S. 2012. Dopamine, affordance and active inference. PLoS Comp Biol, 8(1), e1002327. doi: 10.1371/journal.pcbi.1002327

Gallivan JP, McLean DA, Flanagan JR, Culham JC. 2013. Where one hand meets the other: limb-specific and action-dependent movement plans decoded from preparatory signals in single human frontoparietal areas. 33(5), 1991–2008.

Gallivan JP, McLean DA, Smith FW, Culham JC. 2011. Decoding effector-dependent and effector-independent movement intentions from human parieto-frontal brain activity. J Neurosci, 31(47), 17149–68.

Gentilucci M, Gangitano M. 1998. Influence of automatic word reading on motor control. Eur J Neurosci 10, 752–756.

Gentilucci M. 2002. Object motor representation and reaching-grasping control. Neuropsychologia 40(8): 1139–53.

Gibson JJ. 1979. The ecological approach to visual perception. Boston: Houghton Mifflin

Glasser MF, Coalson TS, Robinson EC, Hacker CD, Harwell J, Yacoub E, et al. 2016. A multi-modal parcellation of human cerebral cortex. Nature 536(7615), 171–178. doi: 10.1038/nature18933.

Glover S, Miall RC, Rushworth MF. 2005. Parietal rTMS disrupts the initiation but not the execution of on-line adjustments to a perturbation of object size. J Cogn Neurosci 17(1), 124–36.

Gomez MA, Skiba RM, Snow JC. 2017. Graspable objects grab attention more than images do. Psychol Science doi.org/10.1177/0956797617730599

Goodale, MA, Milner, AD. 1992. Separate visual pathways for perception and action. Trends Neurosci 15(1), 20–25.

Grafton, ST, Hamilton, AF. 2007. Evidence for a distributed hierarchy of action representation in the brain. Hum Mov Sci 26(4), 590–616. doi: 10.1016/j.humov.2007.05.009.

Grezes J, Tucker, M., Armony, J, Ellis, R, Passingham, RE. 2003. Objects automatically potentiate action: an fMRI study of implicit processing. Eur J of Neurosci 17, 2735–2740.

Grol MJ, Majdandzic J, Stephan KE, Verhagen L, Dijkerman HC, Bekkering H, Verstraten FAJ, Toni I. 2007. Parieto-frontal connectivity during visually guided grasping. J Neurosci 27 (44), 11877–11887.

Harris, CM, Wolpert, DM. 1998. Signal-dependent noise determines motor planning. Nature 394(6695), 780–784. doi: 10.1038/29528.

Herbort, O, Butz, MV. 2011. Habitual and goal-directed factors in (everyday) object handling. Exp Brain Res 213(4), 371–382. doi: 10.1007/s00221-011-2787-8.

Holler DE, Fabbri S, Snow JC. 2020. Objecct responses are highly malleable, rather than invariant, with changes in object appearance. Scientific Reports 10: 4654 doi.org/10.1038/s41598-020-61447-8.

Jeannerod, M. 1994. The representing brain: Neural correlates of motor intention and imagery. Behav Brain Sci 17(2), 187–202. doi: 10.1017/S0140525X00034026.

Kornblum S., Lee J. W. 1995. Simulus-response compatibiliy with relevant and irrelevant stimulus dimensions that do and do not overlap with response. Journal of Experimental Psychology: Human Perception and Performance, 21, 855–875.

Kumar S, Yoon EY, Humphreys GW. 2012. Perceptual and motor-based responses to hand actions on objects: evidence from ERPs. Exp Brain Res 220(2), 153–64.

Lee C-I, Mirman D, Buxbaum LJ. 2014. Abnormal dynamics of activation of object use information in apraxia: evidence from eyetracking. Neuropsychologia 59: 13–26

Lewis JW. 2006. Cortical networks related to human use of tools. The Neuroscientist 12(3): 211–231.

Lingnau A, Downing PE. 2015. The lateral occipitotemporal cortex in action. Trends Cogn Sci 19(5), 268–77. doi: 10.1016/j.tics.2015.03.006.

Mahon BZ, Milleville SC, Negri GAL, Rumiati RI, Caramazza A, Martin A. 2007. Action-related properties of objects shape object representations in the ventral stream. Neuron 55(3): 507–520.

Makris S, Hadar AA, Yarrow K. 2011. Viewing objects and planning actions: on the potentiation of grasping behaviours by visual objects. Brain Cogn 77(2), 257–64.

McBride, J, Sumner, P, Husain, M. 2012. Conflict in object affordance revealed by grip force. Q J Exp Psychol 65(1), 13–24. doi: 10.1080/17470218.2011.588336.

Monaco S, Chen Y, Medendorp WP, Crawford JD, Fiehler K, Henriques DYP. 2014. Functional Magnetic Resonance Imaging Adaptation Reveals the Cortical Networks for Processing Grasp-Relevant Object Properties. Cereb Cortex 24:1540–1554.

Nachev P, Kennard C, Husain M. 2008. Functional role of the supplementary and pre-supplementary motor areas. Nat Rev Neurosci 9(11): 856–69.

Owen AM. 1997. Cognitive planning in humans: neuropsychological, neuroanatomical and neuropharmacological perspectives. Prog Neurobiol 53(4), 431–50.

Packard MG, Knowlton BJ. 2002. Learning and memory functions of the basal ganglia. Ann Rev Neurosci 25: 563–93.

Petrides M. 2018 Atlas of the morphology of the human cerebral cortex on the average MNI brain. 1^st^ Ed. Academic Press.

Phillips J.C., W. R. 2002. S-R correspondence effects of irrelevant visual affordance: Time course and specificity of response activation. Visual Cognition, 9, 540–558.

Pizzamiglio G, Zhang Z, Duta M, Rounis E. 2020 Factors influencing manipulation of a familiar object in patients with limb apraxia after stroke. Front Hum Neurosci, 13, 465, doi: 10.3389/fnhum.2019.00465.

Praamstra P, Kleine BU, Schnitzler A. 1999. Magnetic stimulation of the dorsal premotor cortex modulates the Simon effect. Neuroreport 10, 3671–3674.

Rice NJ, Valyear KF Goodale MA, Milner AD, Culham JC. 2007. Orientation sensitivity to graspable objects: an fMRI adaptation study. Neuroimage 36(Supp 2), T87–93.

Riddoch, MJ, Edwards, MG, Humphreys, GW, West, R, Heafield, T. 1998. Visual affordances direct action: neuropsychological evidence from manual interference. Cogn Neuropsychol 15(6-8), 645–683. doi: 10.1080/026432998381041.

Rizzolatti, G, Matelli, M. 2003. Two different streams form the dorsal visual system: anatomy and functions. Exp Brain Res 153(2), 146–157. doi: 10.1007/s00221-003-1588-0.

Rosenbaum, DA, Marchak, F, Barnes, HJ, Vaughan, J, Slotta, JD, Jorgensen, MJ. 1990. “Constraints for action selection: Overhand versus underhand grips,” in Attention and performance 13: Motor representation and control. (Hillsdale, NJ, US: Lawrence Erlbaum Associates, Inc), 321–342.

Rosenbaum, DA, Vaughan, J, Barnes, HJ, Jorgensen, MJ. 1992. Time course of movement planning: selection of handgrips for object manipulation. J Exp Psychol Learn Mem Cogn 18(5), 1058–1073.

Rosenbaum, DA, Cohen, RG, Meulenbroek, RGJ, Vaughan, J. 2006. “Plans for Grasping Objects,” in Motor Control and Learning, eds. M.L. Latash & F. Lestienne. (Boston, MA: Springer US), 9–25.

Rothi LJG, Ochipa C, Heilman KM. 1991. A cognitive neuropsychological model of praxis. Cogn Neuropsych 8, 443–458.

Rounis E, Yarrow K, Rothwell JC. 2007. Effects of rTMS conditioning over the fronto-parietal network on motor versus visual attention. J Cogn Neurosci 19(3), 513–524. doi: 10.1162/jocn.2007.19.3.513.

Rounis E, Humphreys G. 2015. Limb apraxia and the “affordance competition hypothesis”. Front Hum Neurosci 9, 429. doi: 10.3389/fnhum.2015.00429.

Rounis E, Zhang Z, Pizzamiglio G, Duta M, Humphreys GW. 2017. Factors influencing planning of a familiar grasp to an object: what it is to pick a cup. Exp Brain Res 235(4), 1281–1296. doi: 10.1007/s00221-017-4883-x.

Rounis E, van Polanen V, Davare M. 2018. A direct effect of perception on action when grasping a cup. Sci Rep 8(1): 171. doi: 10.1038/s41598-017-18591-5.

Rushworth MF, Nixon PD, Renowden S, Wade DT, Passingham RE. 1997. The left parietal cortex and motor attention. Neuropsychologia 35(9): 1261–73.

Rushworth MF, Ellison A, Walsh V. 2001. Complementary localization and lateralization of orienting and motor attention. Nat Neurosci 4(6), 656–661. doi: 10.1038/88492.

Sakreida K, Effnert I, Thill S, Menz MM, Jirak D, Eickhoff CR, Ziemke T, Eickhoff SB, Borghi AM, Binkofski F. 2016. Affordance processing in segregated parieto-frontal dorsal stream sub-pathways. Neurosci Biobehav Rev 69, 89–112. doi: 10.1016/j.neubiorev.2016.07.032.

Simon, J. R. 1969. Reaction toward the source of stimulation. Journal of Experimental Psychology, 81, 174–176

Sirigu A, Cohen L, Duhamel JR, Pillon B, Dubois B, Agid Y. 1995. A selective impairment of hand posture for object utilization in apraxia. Cortex 31(1), 41–55.

Snow JC, Pettypiece CE, McAdam TD, McLean AD, Stroman PW, Goodale MA, Culham JC. 2011. Bringing the real world into the fMRI scanner: repetition effects for pictures versus real objects. Sci Rep 1: 130. Doi: 10.1038/srep00130.

Till BC, Masson MEJ, Bub DN, Driessen PF. 2014. Embodied effects of conceptual knowledge continuously perturb the hand in flight. Psychol Science 25(8), 1637–1648.

Tucker M, Ellis R. 1998. On the relations between seen objects and components of potential actions. J Exp Psychol Hum Percept Perform 24(3), 830–846. doi: 10.1037//0096-1523.24.3.830.

Tucker M, Ellis R. 2001. The potentiation of grasp types during visual object categorisation. Vis Cogn 8, 769–800.

Tunik E, Frey SH, Grafton ST. 2005. Virtual lesions of the anterior intraparietal area disrupt goal-dependent on-line adjustment of grasp. Nat Neurosci 8, 505–511.

Valyear KF, Culham JC, Sharif N, Westwood D, Goodale MA. 2006. A double dissociation between sensitivity to changes in object identity and object orientation in the ventral and dorsal visual streams: a human fMRI study. Neuropsychologia 44(2): 218–28.

Valyear KF, Cavina-Pratesi C, Stiglick AJ, Culham JC. 2007. Does tool-related fMRI activity within the intraparietal sulcus reflect the plan to grasp? Neuroimage 36(Supp 2): T94–T108.

Valyear KF, Gallivan JP, McLean DA, Culham JC. 2012. fMRI repetition suppression for familiar but not arbitrary actions with tools. J Neurosci 32(12): 42247–59.

van Elk M, van Schie H, Bekkering H. 2014. Action semantics: a unifying conceptual framework for the selective use of multimodal and modality-specific object knowledge. Phys Life Rev 11, 220–50.

van Polanen V, Davare M. 2015. Interactions between dorsal and ventral streams for controlling skilled grasp. Neuropsychologia 79(Pt B), 186–191. doi: 10.1016/j.neuropsychologia.2015.07.010.

Ward NS, Bestmann S, Hartwigsen G, Weiss MM, Christensen LOD, Frackowiak RSJ, Rothwell JC, Siebner HR. 2010. Low-frequency transcranial magnetic stimulation over left dorsal premotor cortex improves the dynamic control of visuospatially cued actions. J Neurosci 30(27), 9216–9223.

Watson CE Buxbaum LJ. 2014. Uncovering the architecture of action semantics. J Exp Psychol Hum Percept Perform 40(5): 1832–1848.

Watson CE, Buxbaum LJ. 2015. A distributed network critical for selecting among tool-directed actions. Cortex 65: 65–82.

Verhagen L, Dijkerman HC, Grol MJ, Toni I. 2008. Perceptuo-motor interactions during prehension movements. J Neurosci 28:4726–4735

Verhagen L, Dijkerman HC, Medendorp WP, Toni I. 2012. Cortical dynamics of sensorimotor integration during grasp planning. J Neurosci 32:4508–4519

Vesia M, Prime SL, Yan X, Sergio LE, Crawford JD. 2010. Specificity of Human Parietal Saccade and Reach Regions during Transcranial Magnetic Stimulation. J Neurosci 30:13053–13065.

Waszak F, Wascher E, Keller P, Koch I, Aschersleben G, Ronsenbaum DA, Prinz W. 2005. Intentionbased and stimulus-based mechanisms in action selection. Exp Brain Res 162: 346–356. doi:10.1007/s00221-004-2183-8.

Wilf M, Holmes NP, Schwartz I, Makin TR. 2013. Dissociating between object affordances and spatial compatibility effects using early response components. Front Psychol 4: 591. doi: 10.3389/fpsyg.2013.00591.

Wolpert DM, Miall RC. 1996. Forward Models for Physiological Motor Control. Neural Netw 9(8), 1265–1279. doi: 10.1016/s0893-6080(96)00035-4.

Wolpe N, Hezemans FH, Rowe JB. 2020. Alien limb syndrome: a Bayseian account of unwanted actions. Cortex 127: 29–41.

Wolpert DM. 1997. Computational approaches to motor control. Trends Cogn Sci, 1(6), 209–216.

Wong, AL, Haith, AM, Krakauer, JW. 2015. Motor Planning. Neuroscientist 21(4), 385–398. doi: 10.1177/1073858414541484.

Wurm MF, Caramazza A, Lingnau A. 2017. Action categories in lateral occipitotemporal cortex are organised along sociality and transitivity. J Neurosci, 37(3), 562–575.

Xia M, Wang J, He Y. 2013. BrainNet Viewer: A Network Visualization Tool for Human Brain Connectomics. PLoS ONE 8: e68910.

Zimmermann M, Toni I, de Lange FP. 2013. Body posture modulates action perception. J Neurosci 33(14), 5930–5938.

Zimmermann M, Mars RB, de Lange FP, Toni I, Verhagen L. 2018. Is the extrastriate body area part of the dorsal visuomotor stream? Brain Struct Funct 223, 31–46.

